# A New Straightforward Method for Automated Segmentation of Trabecular Bone from Cortical Bone in Diverse and Challenging Morphologies

**DOI:** 10.1101/2021.03.02.433409

**Authors:** Eva C. Herbst, Alessandro A. Felder, Lucinda A. E. Evans, Sara Ajami, Behzad Javaheri, Andrew A. Pitsillides

## Abstract

Many physiological, biomechanical, evolutionary and clinical studies that explore skeletal structure and function require successful separation of trabecular from cortical compartments of a bone that has been imaged by X-ray micro-computed tomography (microCT) prior to analysis. Separation is often time-consuming, involves user bias and needs manual sub-division of these two similarly radio-opaque compartments. We have developed an objective, automated protocol which reduces user bias and enables straightforward, user-friendly segmentation of trabecular from cortical bone without requiring sophisticated programming expertise. This method can conveniently be used as a “recipe” in commercial programmes (Avizo herein) and applied to a variety of datasets. Here, we characterise and share this recipe, and demonstrate its application to a range of murine and human bone types, including normal and osteoarthritic specimens, and bones with distinct embryonic origins and spanning a range of ages. We validate the method by testing inter-user bias during the scan preparation steps and confirm utility in the architecturally challenging analysis of growing murine epiphyses. We also report details of the recipe, so that other groups can readily re-create a similar method in open access programs. Our aim is that this method will be adopted widely to create a more standardized and time efficient method of segmenting trabecular and cortical bone.

## 1. Introduction

Biomechanists, bone physiologists, biologists, clinicians, and palaeontologists analyse bone structure to answer a myriad of questions (Bishop et al. 2018, Hoechel et al. 2015, Reznikov et al. 2020, Rueda et al. 2006). Some whole bone analyses rely on the presumed conservation of development, remodelling and repair factors across cortical and trabecular bone compartments, yet most studies investigate these two bone types separately due to the likelihood that inherent differences in their behavior and responsiveness exist (Lavigne et al. 2005; Simon et al. 2008; Wade-Gueye et al. 2010). Compared to the cortex, trabecular bone can constitute a small fraction of total volume, is more porous and less dense, and via its large surface area for remodelling supplies most of the exchangeable calcium pool (Aerssens et al. 1997). This illustrates the differing functions and performance of these structurally diverse compartments, which have been reported to extend to bone type-related differences in osteoblast behaviour even when isolated and maintained in vitro (Shah et al. 2015). Due to these differences, it is paramount that most studies investigate the cortical or trabecular compartments in isolation, which necessitates the effective segmentation of these two types of bone.

For computed tomography (CT) data, a manual pre-processing segmentation step is commonly used to differentiate cortical and trabecular bone (Florea et al. 2015, Gohin et al. 2020, Javaheri et al. 2018, Poulet et al. 2016, Whitmarsh et al. 2019), but is both time-intensive and subjective. In particular, it is difficult to determine where the intersection between the trabeculae and cortical bone starts. In a 3D stack of images, the *base* of a trabecular column could be characterised either as a trabecular or cortical component, depending on which slice is viewed. The guidelines for defining trabecular boundaries in manual segmentations are rarely reported (Gohin et al. 2020, Poulet et al. 2016, Javaheri et al. 2018; Liu et al. 2010). Another commonly used alternative to manual segmentation is to crop a region of interest (for example a sphere or cube) from the trabecular region and analyse only this volume (Bishop 2018, Doube et al. 2011, Hoechel et a. 2015, Lui et al. 2010). However, this approach excludes data from the trabecular regions outside of this volume, making it likely that changes near the cortical boundary would therefore be missed.

Previous studies have also developed automated segmentation protocols, but they have various limitations or are time consuming to implement. Methods avoiding manual segmentation of 2D slices use bimodal methods that rely on single grey value thresholding, which tend to fail in complex scenarios (Spoor et al. 1993) or gradient-based edge detection algorithms which perform well (Scherf and Tilgner 2009, Zebaze et al. 2013). For example, Lublinsky et al. (2007) built a 5-step freely-available algorithm that mapped the periosteal edge to create a cortical mask, the innermost edge of which defined the cortical: trabecular interface. This however requires specific definition of filter, threshold, and categorization of cortical thickness (thick or thin) prior to analysis, and that the volume of the space outside the bone exceeds the volume of the inter-trabecular space, thus restricting its utility to only some CT images. Some segmentation methods require constant thickness of the cortical bone (Treece et al. 2010, Whitmarsh et al. 2019), an assumption which does not apply to many bones, or require that the thickness of the cortical bone is greater than that of the trabeculae and that the cortical border is continuous (Ang et al. 2019). Other methods require a template (reference segmentation) or even entire training set of segmented images that encompass the range of variation expected in the sample set (Rueda et al. 2006, Väänänen et al. 2019). Creating such a training set would require time spent on manual segmentation of the initial reference samples. Furthermore, for fossil specimens, there may not be enough material present to create any reference samples.

The methods by Buie et al. 2007 and Burghardt et al. 2010 (which builds on the former) solve many of these issues. These methods are, however, designed for specific morphologies and require certain conditions about the bone distribution to be met, specifically relating to the diameter (in pixels) of the largest pore (or sinus) connecting the marrow to the exterior. Such large sinuses require extensive amounts of dilation and erosion to *close* the boundaries of the regions of interest in order for a connectivity filter to effectively distinguish marrow from outside space, but large amounts of dilation and erosion will result in excessive smoothing of the cortical-trabecular boundary. To address these limitations, we have developed an objective, automated protocol for segmenting trabecular and cortical bone which was validated in animal and human bone samples. We specifically wanted to develop a method that was automated but also flexible enough to draw on the user’s anatomy knowledge to define the marrow space across various challenging morphologies. Our method relies on an input segmentation of the marrow space and can easily be loaded as a “recipe” in Avizo, which enables all further steps to run automatically. We compare our method to other automatic segmentation methods; this comparison is not intended to be exhaustive but rather a discussion of the range of methods available and how our method builds upon and differs from these.

## 2. Materials and Methods

### 2.1 Micro-CT Datasets

The automated segmentation algorithm was initially evaluated in knee joint epiphyses of normal healthy control (CBA) and osteoarthritis-prone (STR/Ort) mouse strains, followed by samples from the human femoral head, skull and vertebrae. The micro-CT dataset originated from: (1) tibial epiphysis of a skeletally-mature 20-week-old normal CBA mouse (doi.org/10.6084/m9.figshare.14097380), (2) tibial epiphysis of a 19-week-old osteoarthritis-prone STR/Ort mouse (doi.org/10.6084/m9.figshare.14097983), (3) tibial epiphysis of an ageing 51-week-old osteoarthritis-prone STR/Ort mouse (doi.org/10.6084/m9.figshare.14098052), (4) part of a human femoral head from a 77-year-old female (doi.org/10.6084/m9.figshare.14097263), (5) human vertebra from 24-year-old female (data from Mills and Boyde 2020), (6) human parietal bone from a 5-month-old male with craniosynostosis (doi.org/10.6084/m9.figshare.14135879), and (7) tibial epiphysis of a young, growing 8-week-old STR/Ort mouse (doi.org/10.6084/m9.figshare.14141159). The latter two specimens were chosen specifically for the high porosity and sinus content of their developing primary cortical bone. The voxel size of the mouse CT scans was 0.005 mm, the voxel size of the human vertebra scan was 0.03 mm, the voxel size of the human skull scan was 0.009 mm, and the voxel size of the human femur scan was 0.01 mm. The human samples were retrieved following ethical approval and patient consent. Ethical approval for the animal procedures was carried out in accordance with the Animals (Scientific Procedures) Act 1986, an Act of Parliament of the United Kingdom, approved by the Royal Veterinary College Ethical Review Committee and the United Kingdom Government Home Office. Our CT scan data is available in the following Figshare repository: https://figshare.com/projects/Trabecular_and_Cortical_Bone_Segmentation_Method/99434

### 2.2 Application of Algorithm to the 3D Dataset

The datasets first required some semi-automated preprocessing to produce the marrow space segmentation. The marrow space segmentation is then used as the input for the trabecular segmentation algorithm. The preprocessing of CT data and implementation of the algorithm was performed in Avizo (Thermo-Scientific, v.2019.2 and 2020.1).

#### 2.2.1 Preprocessing of Dataset

We preprocessed the dataset in Avizo, although other segmentation software could also be used. First, we filtered the CT scans using a non-local means filter (3D GPU adaptive manifold setting). Next, we separated the bone of interest from the rest of the scan by placing “seeds” and assigning them to two different regions (defined as “materials” in Avizo): the first containing both the bone of interest and its marrow space; and the second being the background (any bone(s) not included in analysis and background voxels). Then a watershed operation was performed on these “seeds’’. The watershed algorithm assigns regions based upon the positioning of these different seeds, using changes in gradient of the voxel greyscale values to determine the boundaries of these regions. For the murine samples, the bone of interest was the entire tibial epiphysis, which was removed from the metaphysis by the watershed operation. For the human samples, all bone present in the scan was included in the analysis. This enables a semi-automated segmentation of the whole region of interest (mouse epiphysis in Fig. 1B) that is quicker and less biased than manual segmentation.

**Fig. 1.**
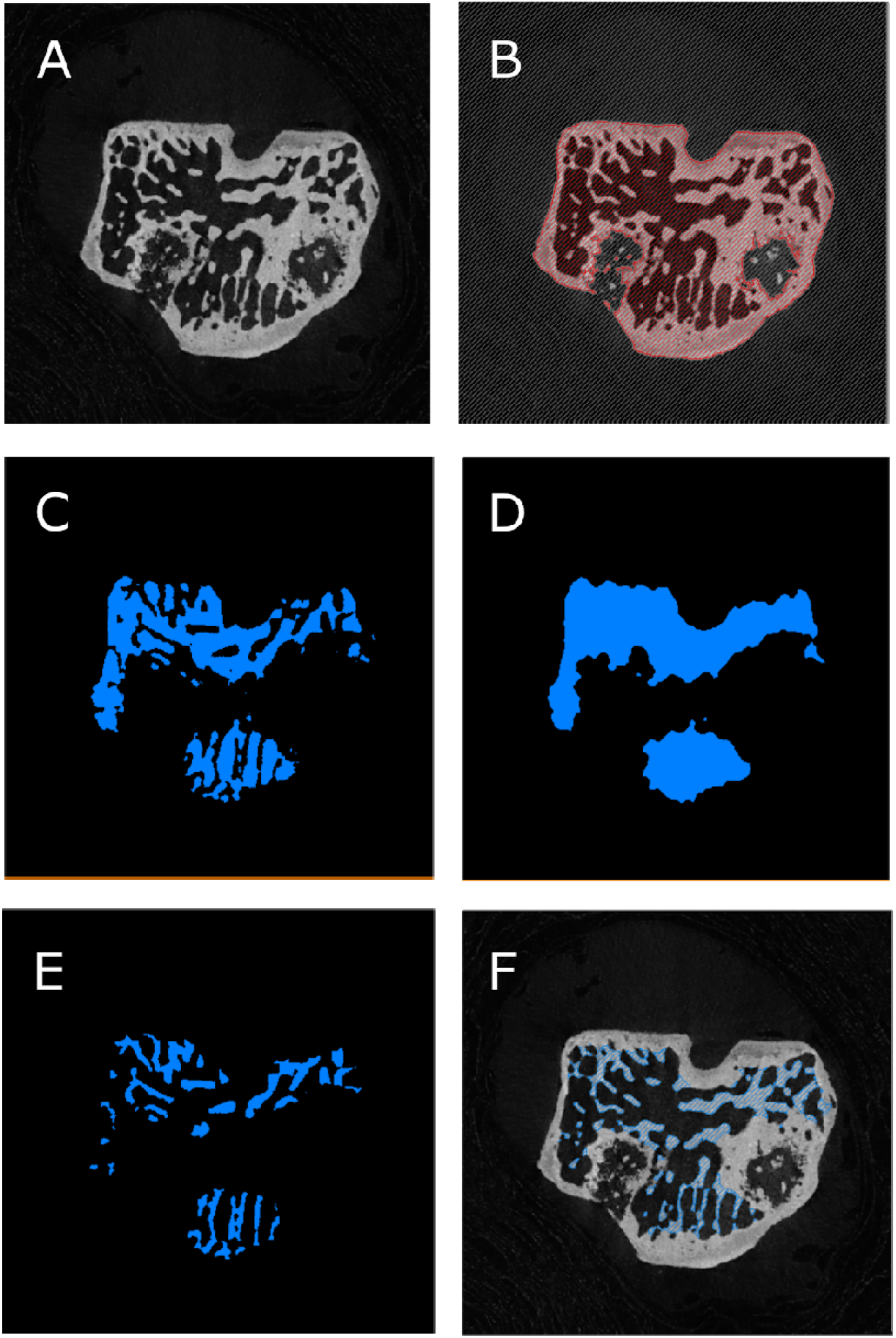
Preparation and step-wise application of algorithm, demonstrated within the tibial epiphysis of a 19-week-old STR/Ort mouse. A) Filtered CT scan of mouse epiphysis; B) epiphysis (both bone and marrow space) selected via the watershed operation; C) the marrow space isolated by thresholding of greyscale values; D) the marrow space is shrink-wrapped via the closing algorithm; E) the trabeculae are isolated by subtracting the marrow space (C) from the shrink-wrapped space (D); F) the resulting trabecular segmentation shown on the original scan.

Growth plate bone bridges (radiopaque tissue connecting the metaphysis and the epiphysis, see Staines et al. 2018) were manually removed from the tibial epiphysis. For the human samples, the bone and marrow space were separated from the background using the same approach, although the growth bridge removal was not required. The scans of the human bone samples were therefore much quicker to pre-process. This preprocessing was followed by the use of appropriate thresholding ranges to separate the marrow space from the bone (greyscale values of 0-70 for the mouse scans, 0-37 for the human femoral head, 0-30 for the human vertebra, and 0-49 for the human skull) to segment out the marrow space. The marrow space is the input for the automatic method.

#### 2.2.2 Segmentation of Trabecular Bone

The algorithm consists of several sequential steps that were applied to a dataset (Fig.1A) after isolating the marrow space (Fig. 1B, see preprocessing, section 2.2.1). Essentially, the algorithm starts by smoothing the input marrow space segmentation (Fig. 1C), shrink-wrapping the smoothed marrow space (Fig. 1D), reducing noise by another smoothing operation, and finally selecting the bone inside the smoothed shrink-wrapped volume. This bone corresponds to the trabecular compartment (Fig. 1E,F). The first smoothing step is achieved by a 3 pixel ball erosion and a 3 pixel ball dilation. This step removes the small sinuses connecting the marrow space to the exterior of the bone; these need to be removed so that they are not included in the subsequent shrink wrapping step. The “shrink wrapping” is achieved by a ball-closing operation with a value of 25. This value was chosen based on iterative tests on the mouse samples, and worked well for the human samples. The closing operation takes a binary image and fills small holes, smoothes object boundaries, and connects close objects to produce a new binary image. The second smoothing step is achieved by a 1 pixel ball erosion and 1 pixel ball dilation to reduce noise and remove aberrant single pixels. This algorithm was implemented as an Avizo recipe and applied to the micro-CT datasets described above.

### 2.3 Method for Addressing the Presence of Intra-cortical Sinuses and Porosity

We tested the effect of including small sinuses that connect the marrow space to the epiphyseal exterior before applying the algorithm. These sinuses are usually included when the watershed operation segments the bone together with the associated marrow space, from the surrounding background (Figure 2A). When these sinuses are included in the marrow space, the algorithm shrink wraps the sinus and therefore inappropriately designates adjacent cortical bone as trabecular bone (Figure 2B). This can be avoided if sinuses are excluded from the inter-trabecular space layer, by manually removing the sinus to the extent of a boundary drawn through the main marrow cavity in the adjacent slices (Figure 2C,D). Note that very small sinuses (up to 3 pixel width) are removed in a step in the automatic algorithm (Figure 2A).

**Fig. 2.**
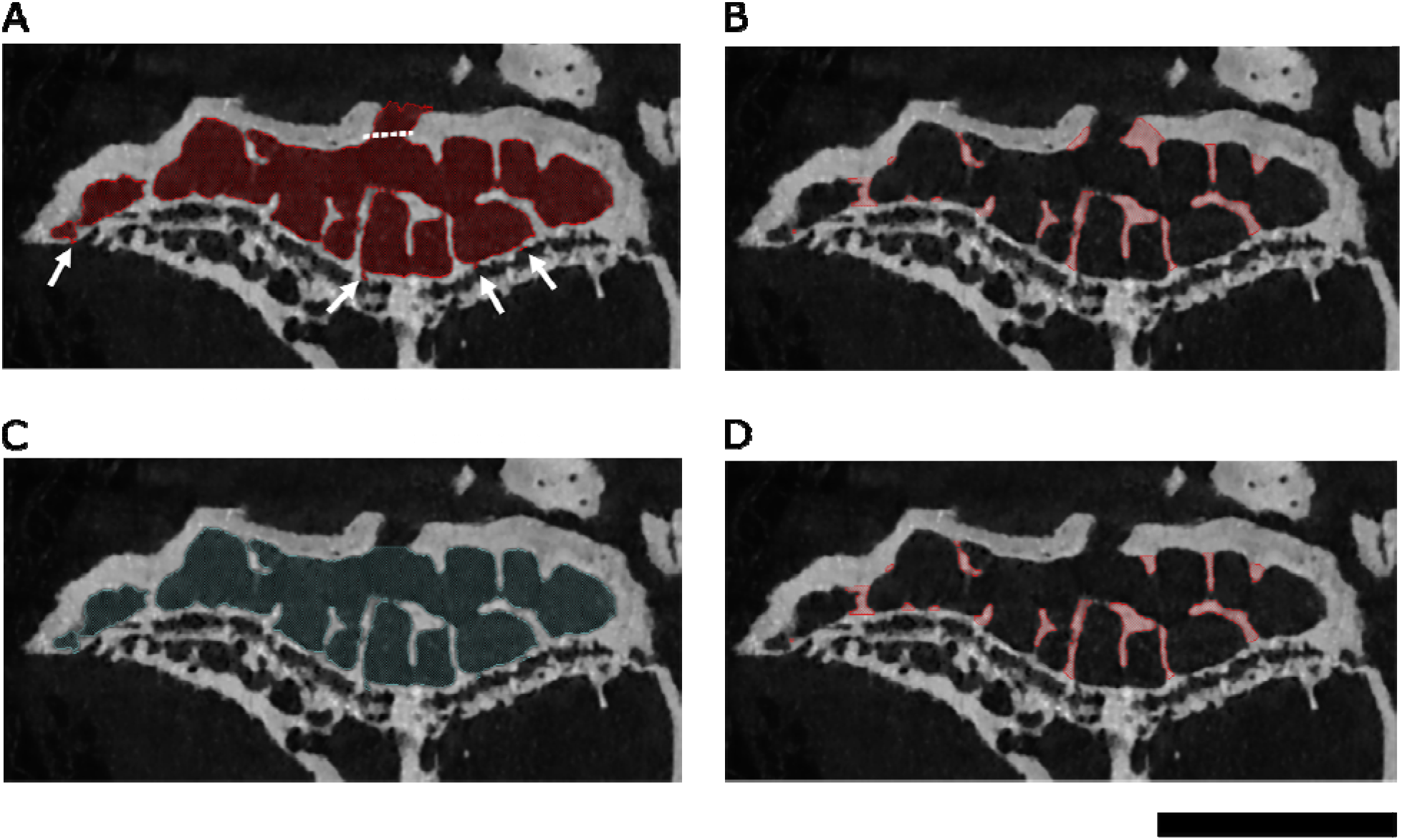
Sinus cleaning demonstrated in the tibial epiphysis of a 51-week-old STR/Ort mouse. A) shows the marrow space without sinus cleaning, and the resulting trabeculae are shown in B). In C), the marrow space was manually cleaned (along the dotted line shown in A). D) shows the result of the cleaned marrow space. Very small sinuses, such as those shown by the arrows in A), are automatically removed by the algorithm to prevent them from being shrink-wrapped. Scale bar = 1 mm.

We ran the algorithm on sinus-cleaned and non-cleaned versions of both a 19-week-old and a 51-week-old STR/Ort mouse tibial epiphysis. To assess the effect of the cleaning step, we compared the relative difference (%) between the two versions for the following morphological parameters: trabecular BV/TV, trabecular volume/cortical volume, and anisotropy. Relative difference was calculated as the difference between the values obtained from the sinus-cleaned and sinus-uncleaned versions, divided by the sinus-cleaned version. Percentages were rounded to the nearest 0.01 percent. We anticipated that in larger bones (such as human vertebrae and limb bones), the effect of sinus cleaning would have even less impact on results, since sinuses in these bones will be much smaller relative to the total bone volume. To test this hypothesis, we also analysed part of a human vertebra (24 year old female, data from Mills and Boyde 2020) and compared outputs from two segmentations, including and excluding sinus cleaning respectively. We compared the relative difference (%) for the same parameters as described above for the mouse samples.

To determine any potential effects of increased cortical porosity on the functionality of our recipe, we examined the outcome when the recipe was applied to the human parietal bone of a 5-month-old male and the tibial epiphysis of an 8-week-old STR/Ort mouse; the cortical bone compartment of these immature bones is particularly porous. Cortical porosity in the human sample consists largely of growing primary cortical bone, with non-consolidated osteons in the very early stages of in-filling. In the mouse, the thick porous layer of cortical bone could also be interpreted as a much thinner layer of cortical bone with thick trabeculae; a potential source of high inter-user variation in studies lacking an objective segmentation method. We used these samples to test how our algorithm draws the boundary for such specimens with high porosity. We also used these samples to test some manual adjustments (sinus cleaning), to determine how this affects the boundary definition between trabecular and cortical bone.

### 2.4 Validation: Testing Inter-User Variation

Although our algorithm is automatic, some user-input was required in pre-processing the CT images prior to automatic segmentation. An example is the isolation of epiphyses from entire long bones in mouse samples. However, as noted above with regard to sinus cleaning, the benefit of validating the method on such difficult scans is that we anticipate inter-user bias to be even smaller on scans requiring less preparatory manual pre-processing.

To calculate inter-user bias, two experienced users segmented the same scans; one scan of a 19-week-old STR/Ort mouse tibial epiphysis, and one scan of part of a human vertebra (24-year-old female, data from Mills and Boyde 2020) using the automatic algorithm. In the mouse samples, both users tested the method with and without sinus removal. For the human vertebra, both users independently cleaned sinuses. The percent difference between users was determined and the relative difference (%) for the same parameters we used for the sinus cleaning sensitivity tests. Relative difference was calculated as the difference between the values obtained from the two users, divided by the values obtained by user one. As for the sinus cleaning tests, percentages were rounded to the nearest 0.01 percent.

## 3. Results

Our automatic segmentation method performed well on tibial epiphyses of both healthy ageing control (CBA) and osteoarthritis-prone STR/Ort mouse tibial epiphyses at a range of different ages (Fig. 3). The algorithm also successfully segmented cortex from trabeculae in parts of a human vertebra, a human femur, and a human skull bone (Fig. 4). The method successfully segmented trabecular bone from cortical bone for scans with a range of different scan resolutions (0.005 - 0.03 mm). While the preprocessing time varies depending on sinus cleaning and quantity of growth bridges that need to be removed, computation time of the algorithm is consistently quick (under 30 seconds with a 32 GB RAM computer).

**Fig. 3.**
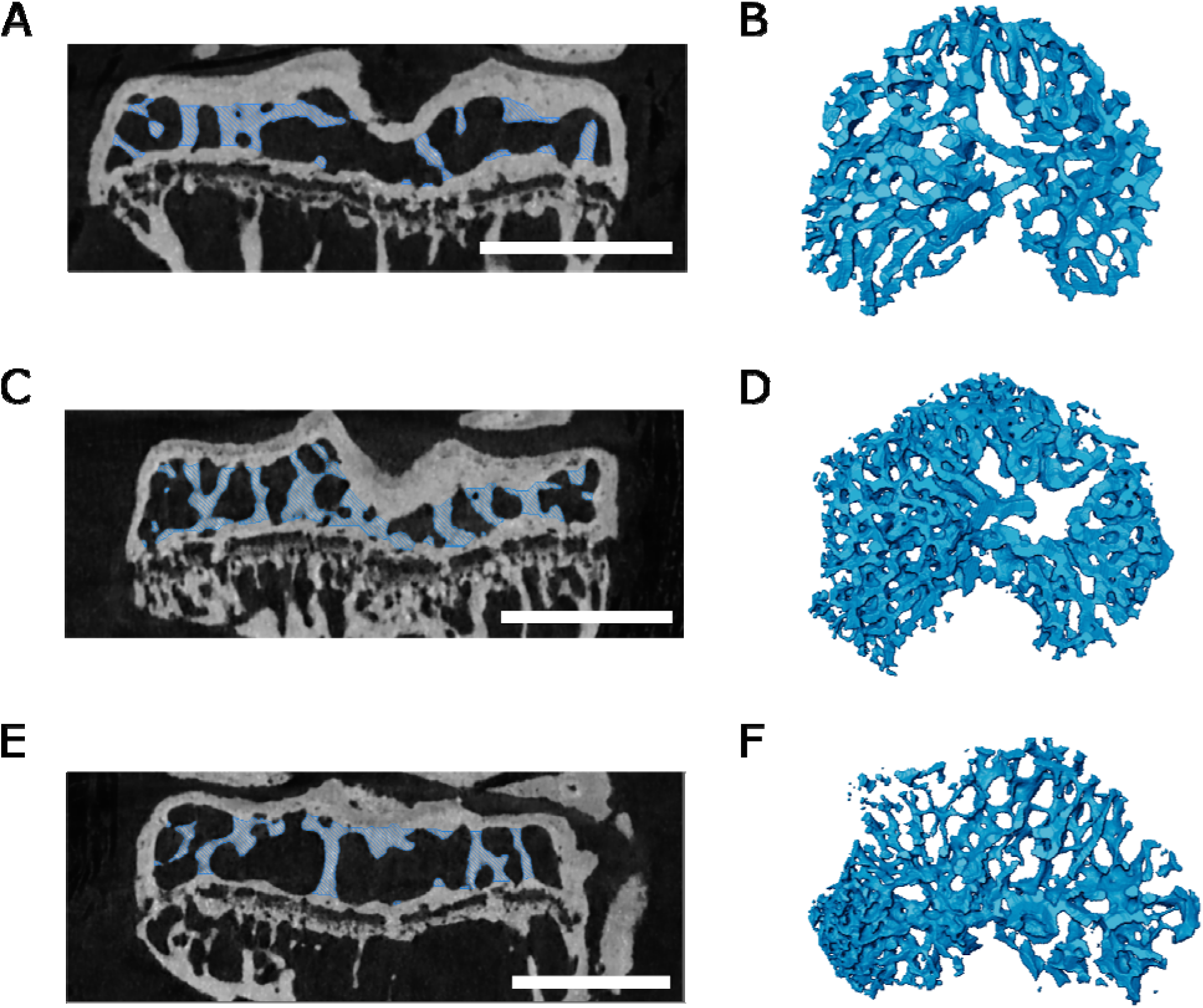
Examples of algorithm applied to mouse tibial epiphyses. A,B) 20-week-old CBA mouse; C,D) 19-week-old osteoarthritic (STR/Ort) mouse; E,F) 51-week-old osteoarthritic (STR/Ort) mouse. A,C,E) CT scan cross sections with trabeculae segmented out; B,D,F) 3D model of segmented trabeculae. Scale bars = 1 mm.

**Fig. 4.**
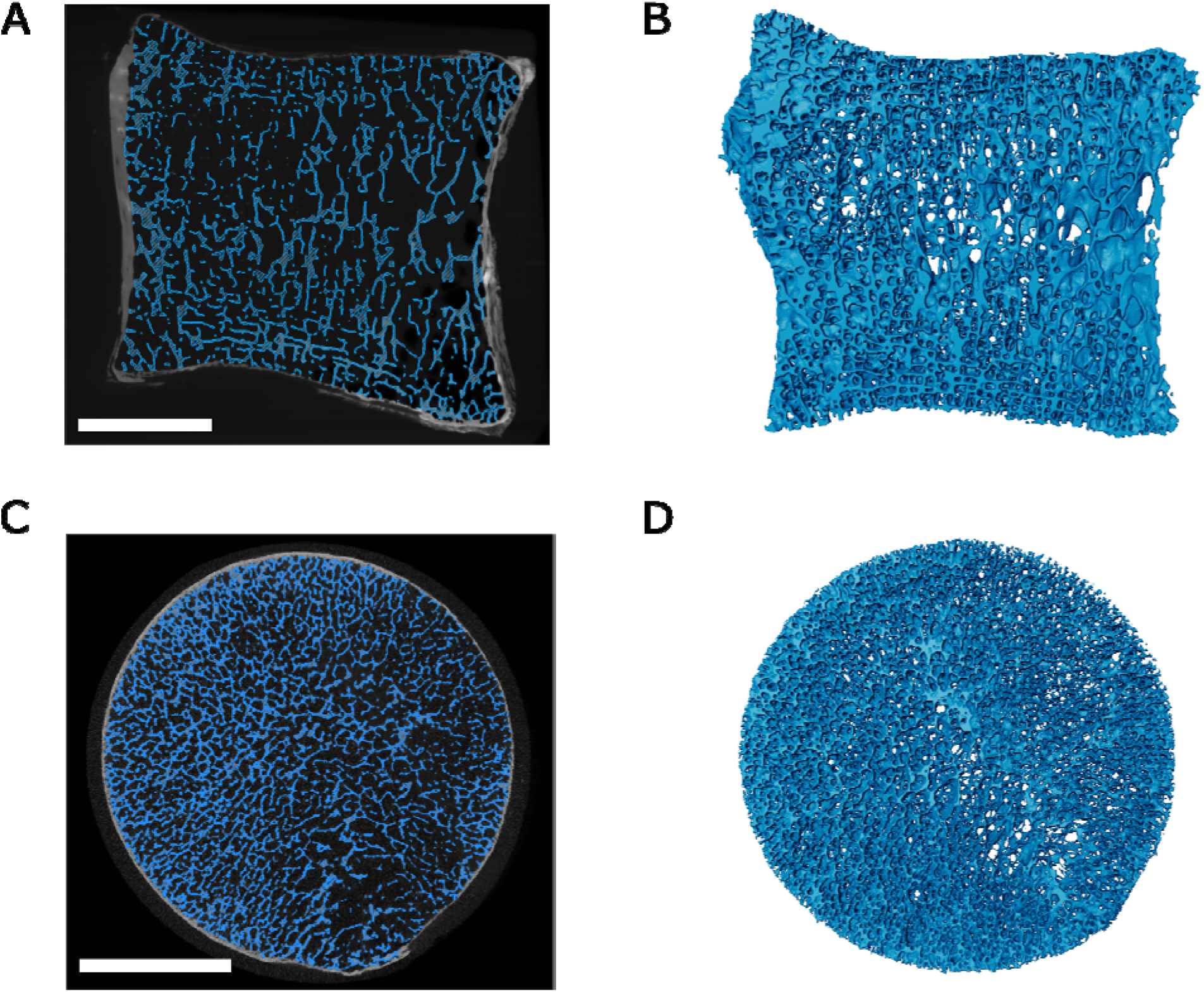
Examples of algorithm applied to human samples. A,B) Part of vertebra from a 24-year-old human female (Mills and Boyde 2020); C,D) Part of human femoral head from a 77-year-old female. A,C) CT scan cross sections with segmented trabeculae highlighted in blue and cortical bone left in greyscale; B,D) 3D model of segmented trabeculae. Scale bars = 10 mm.

### 3.1 Minor Effects of Omitting Sinus Cleaning Step

The difference between samples analysed before and after the sinus cleaning step are reported in Table 1.

**Table 1.**
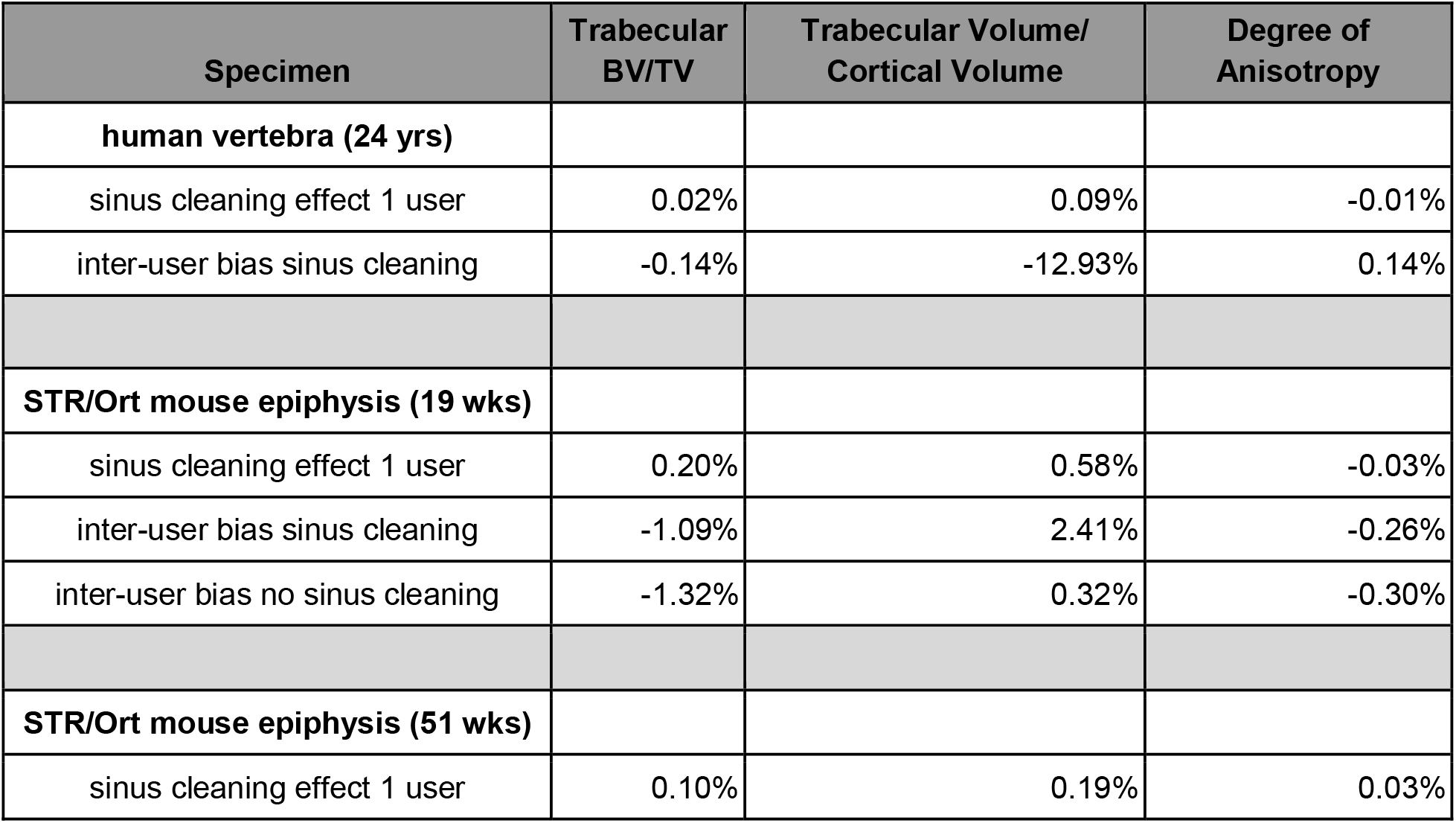
Percentage differences between sinus cleaned and non-sinus cleaned samples, and percentage difference between different users. Percentages rounded to nearest 0.01%.

| Specimen | Trabecular BV/TV | Trabecular Volume/ Cortical Volume | Degree of Anisotropy |
| --- | --- | --- | --- |
| **human vertebra (24 yrs)** |  |  |  |
| sinus cleaning effect 1 user | 0.02% | 0.09% | -0.01% |
| inter-user bias sinus cleaning | -0.14% | -12.93% | 0.14% |
| **STR/Ort mouse epiphysis (19 wks)** |  |  |  |
| sinus cleaning effect 1 user | 0.20% | 0.58% | -0.03% |
| inter-user bias sinus cleaning | -1.09% | 2.41% | -0.26% |
| inter-user bias no sinus cleaning | -1.32% | 0.32% | -0.30% |
| **STR/Ort mouse epiphysis (51 wks)** |  |  |  |
| sinus cleaning effect 1 user | 0.10% | 0.19% | 0.03% |

As predicted, differences between sinus cleaned/non-sinus cleaned results were minimal, and the two versions differed least in the human vertebra where sinus volume is particularly small relative to overall volume. This sensitivity analysis demonstrates that both for the murine epiphyses and human samples, the sinus cleaning step could be omitted with minimum change to results, which would also be expected for sample types with similar sinus volume to total volume ratios.

### 3.2 Specimens with High Porosity

In the human parietal bone, our algorithm first shrink-wrapped a large part of the developing primary cortical bone, classifying solely the most periosteal layer as cortex. This was due to the osteonal pores being connected and of similar size to trabecular pores. To adjust for this, we manually removed the primary osteonal pores from the marrow space segmentation (Fig. 5A,B). In the murine epiphysis (Fig. 5C,D) most sinuses were removed by the algorithm and only a few large sinuses were manually cleaned (see section 2.3 and Figure 2). We tested whether the inclusion of smaller spaces (such as the sinus marked with a red arrow in Figure 5E) affects the trabecular segmentation outcome. There was no effect, because the space was small enough to be automatically removed by the erosion and dilation algorithm steps. Figure 5F shows a close-up of the trabecular/cortical boundary as determined by the algorithm, confirming that the trabecular boundary segmentation is insensitive to small cortical pores.

**Fig. 5.**
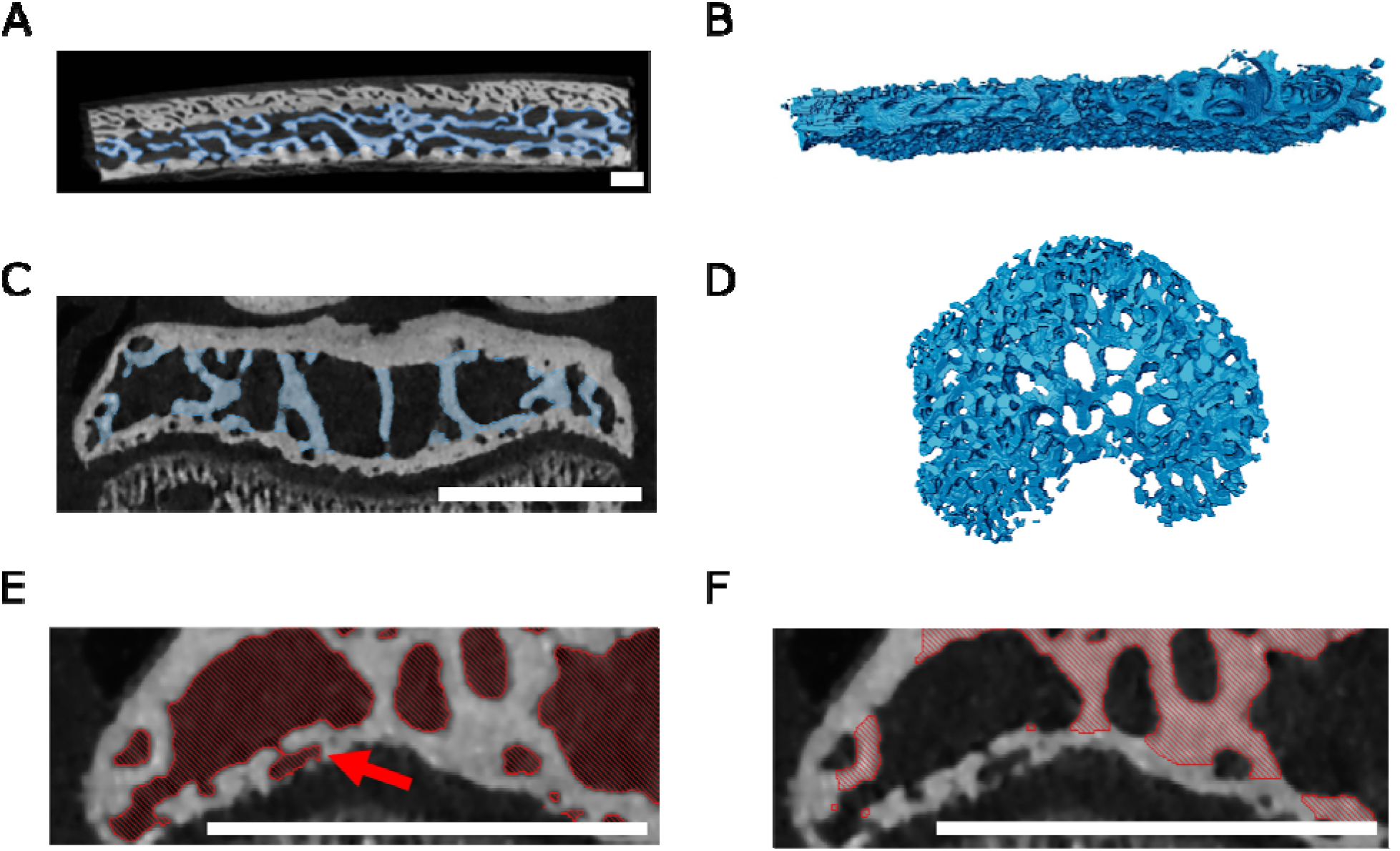
Examples of specimens with high porosity. A,B) Section of human skull bone from a 5-month-old male with craniosynostosis; C,D,E,F) 8-week-old STR/Ort mouse. A,C) CT scan cross sections with trabeculae segmented out; B,D) 3D model of segmented trabeculae. In the human skull, sinuses in the cortical bone were manually removed to ensure that none of the cortical bone was shrink wrapped. In the mouse, including or manually removing the small horizontally oriented space (red arrow) as part of the marrow space (E) did not affect the results (F). This is because this space was not shrink wrapped with the parameters of our algorithm; it was removed during the erosion step. F) close-up of the trabecular segmentation created after running the algorithm on E). Scale bars = 1 mm.

### 3.3 Low Inter-user Bias

Inter-user differences were generally low (see results in Table 1).

## 4. Discussion

Our results demonstrate that this method can be applied to bones from a variety of species, both endochondral and intramembranous origins, and across a range of healthy and pathological specimens. Although this method is not the ‘only’ automatic segmentation method, it is nonetheless of value to our field because it requires no coding background in users, and is therefore accessible and easy to use. Avizo is one of the most commonly used segmentation programs by researchers, and creating an Avizo “recipe” for this method makes it very easy to implement for Avizo users in particular. Loading our recipe in Avizo does not require any coding knowledge; after downloading the “recipe”, the user selects the pre-processed marrow space as the recipe input and all steps will be run in sequence. In addition to making the Avizo “recipe” freely available, we share its specific steps in Supplementary Information so that the method can also be applied in open-source programs which are more accessible and may be preferred by other researchers.

Usefully, our algorithm works on specimens that would not satisfy the assumptions required for application of other automated segmentation methods. For example, Ang et al. (2019) developed a time-efficient segmentation method using thickness ranges to differentiate cortical and trabecular bone; this method would likely overestimate cortical bone in our mouse epiphysis and human vertebra samples, since it would interpret the thick trabeculae adjacent to the cortical bone as actually being cortical bone. Our algorithm is also independent of extra-epiphyseal volume - unlike the automatic algorithm by Lublinsky et al. (2007), which requires that the volume outside exceeds the marrow space volume.

It is relevant to note here that the Buie et al. 2007 method is conceptually very similar to ours. Our “closing” step is similar to their erosion/dilation steps, although we first remove small sinuses by a separate 3 point erosion/dilation of the marrow space, and then apply the closing operation (essentially a larger dilation and erosion step). Our methods differ, however, in not applying dilation and erosion steps on both the cortex and the marrow space to delineate the periosteal and endosteal cortical surfaces, which are then used to create a “mask” to segment the trabeculae. While these steps in the Buie et. al. 2007 method typically enable a more automatic detection of marrow space (using a connectivity filter), it also renders it less flexible. For example, any number of dilation and erosions steps will fail to find a reasonable marrow space in the skull fragment we present herein (Fig 5A, B). This is because either the number of dilations will not be enough to completely disconnect marrow pixels from the outside space, or it will be so high that spaces between trabeculae will be filled completely, causing subsequent erosions to ignore these filled spaces. Burghardt et al. 2010 built on the method of Buie et al. (2007) by further validation, providing a program for implementation (available as an extension of the Scanco scan visualization and analysis software). Future studies could test the differences between eroding and dilating small regions first to remove sinuses and then performing the larger erosion and dilation (as in our method) versus performing one large dilation followed by a large erosion (as in the Buie et al. 2007 and Burghardt et al. 2010 methods).

We, nonetheless, consider both methods valuable and our independent development of a conceptually similar approach highlights the usefulness of all of these methods.

Our method may however reach a different user base, specifically users of Avizo software. Our method also should not cause any clipping issues of the outer cortical boundary. Prior methods (Buie et al. (2007), Burghardt et al. (2010)) were hindered somewhat by the fact that if the space outside of the region of interest is smaller than the dilation and erosion amounts, clipping can occur when the dilation performed to create a “cortical mask” grows beyond the boundaries of the image. Since we only perform the erosion and dilation on the marrow space and not the cortical bone, unlike Buie et al. (2007) and Burghardt (2010), we would not have issues with the clipping of the periosteal boundary of the segmented cortex.

Moreover, we have validated our method on an especially difficult sample, the mouse epiphysis, where the sinuses are very large relative to the total volume. Our method offers the option of straightforward manual input, either by changing the algorithm parameters or manually adjusting the marrow space (for example by cleaning sinuses). This advantage allowed us to deal appropriately with the more complex situations encountered in the human skull and the young mice, where a good marrow space segmentation to input into the algorithm is not straightforward to obtain.

Buie et al. 2007 noted the issues with large Volkmann’s canals and discussed that these need to be closed for the algorithm to work well; this can be solved with a higher amount of erosion and dilation, but this can result in too much smoothing, obscuring small features. The authors proposed a solution in which the periosteal threshold is decreased to create a larger mask - however, these steps require an iterative adjustment of the recipe: iteratively testing to see what erosion and dilation values to apply, whether these fully close the sinuses, and then going back and adjusting the thresholds to prevent any smoothing by large dilations and erosions. Here, we provide an alternative approach by presenting details on how to avoid such smoothing by making manual adjustments to sinuses in the pre-processing steps of our protocol. The method of Burghardt et al. (2010) also enabled manual adjustments (which were required for young subjects with dense trabecular networks and older subjects with endosteal trabecularization). However, in this paper we describe in detail how to conduct the sinus cleaning, and tested the effects of including or excluding these sinuses.

Our method enables the adjustment of the parameters (erosions and dilations and the ball closing operation) to control the size of the sinuses that are excluded automatically (currently all sinuses with a width equal to or less than 3 pixels are removed). This is especially relevant for specimens with lots of cortical porosity, as seen in the young human synostotic bone and the young STR/Ort mouse epiphysis. We also observed lots of cortical porosity in young CBA mice (Fig. 5C,E,F). This cortical porosity could also be interpreted as trabecular bone with a very thin cortical layer, and in these young specimens it is hard to manually define where the cortical bone ends and the trabecular bone begins. Our algorithm does not solve this issue of identification. However, our algorithm does provide a consistent method of determining the cortical: trabecular border that is unbiased, and based on the bone’s morphology (specifically the contours of the marrow space). Our method is based on the steps of the algorithm, and the parameters determining which sinuses are removed via erosion and the closing (shrink wrapping) operation. Studies can adjust these parameters to produce results in agreement with their definition of cortical and trabecular bone. For example, if the experimenter interpreted Fig. 5E as representing a thin layer of cortical bone with thick trabeculae, then they could remove the erosion and dilation steps to broaden the shrink wrapping. Manual input is also possible if the user wants to ensure that certain channels are excluded from the marrow space. This was done for the human skull bone (Fig. 5A,B); see section 3.2. Any such deviations from the standard ‘recipe’ should therefore be highlighted in any publications applying our algorithm or variations thereof, to ensure that adjustments for different samples are appropriately shared.

In our sensitivity analyses of the effect of sinus cleaning, differences between samples with/without sinus cleaning steps were very low; this manual step can be omitted with only minute change (<0.6%) to trabecular BV/TV, trabecular volume/cortical volume, and trabecular degree of anisotropy. Degree of anisotropy was the parameter least affected by sinus cleaning (<0.04% for all samples).

Inter-user differences were generally also very low, again especially for degree of anisotropy. Only one value differed significantly: user variation in trabecular volume/cortical for the partial human vertebra was 12.93%. On closer scrutiny, this was attributable to a calcified ligament attachment site on the outer cortex; one user placed seeds there, including the calcified ligament in the initial region of interest - the other user did not. Other parameters, i.e. trabecular BV/TV anisotropy, were very similar between users. This demonstrates that bias was introduced by user variations in anatomical knowledge, and not the algorithm’s cortical/trabecular separation. Theoretically, this problem could also arise anywhere that density (and thus greyscale values) of mineralized soft tissues are similar to bone densities; particularly entheses. It is recommended that in order to avoid such issues, users should ensure that the objectives of any given study, and anatomy of scanned tissues, are clear to all participants. Seeds determining the desired region of interest (with manual cleaning to remove the mineralized soft tissue if necessary) will then be placed more accurately.

Note that sinus cleaning sometimes increases inter-user variation, but may also decrease the difference between users (relative to the uncleaned version). This is because of variation in the results of the watershed operation - depending on variation in seed placement, the boundary between marrow space and exterior may differ between users or be more similar than the sinus cleaned version. Regardless, the inter-user differences for both sinus cleaned and non-sinus cleaned versions of the mouse specimen were similar and very low in magnitude.

### 4.1 Considerations for Application

We recommend three visual checks: initially, after the watershed that separates the region of interest, then another to establish that marrow space is isolated from large intra-cortical sinuses before running the algorithm, and a third to ensure that the method successfully distinguishes trabecular and cortical bone well on your taxon, element of interest, and scan resolution. Users need to ensure voxel size is not too large relative to structure size. For example, in our study of murine bone the voxel size of CT scans was 0.004997 mm and average trabecular thickness was ~0.06 mm. Our method will not work on scans with a low feature size relative to the voxel size. If each trabecula was ~1-2 voxels, the erosion/dilation (noise removal) algorithm step would erase important morphology. However, flexibility in our method enables users to edit out these noise removal steps if desired, to test suitability for lower quality datasets. Users can also easily adjust the values of the erosion and dilation and closing steps, to fine tune the recipe for their dataset. Note that a few times, there were samples in which the final trabecular bone included a few floating pixels of space as part of the trabecular result. This rare occurrence was in pixels near the shrink wrapping boundary (where the boundary exceeded the original region of interest), for example if a big sinus was not cleaned. This was readily remedied by thresholding the trabecular layer to include only bone (using the same bone - space greyscale threshold initially used to isolate the marrow space).

## 5. Conclusion

Our algorithm is a quick, repeatable method of segmenting trabecular and cortical bone. It can easily be implemented in Avizo, and can also be adopted in open-source programs such as ImageJ. Our algorithm was rigorously tested on a variety of specimens. It does not need any initial segmented reference images, making it ideal for fossil work or for starting on a new project where no training set for the specimens of interest exists. Our method also stands out against most others because it does not require certain assumptions to be met, such as cortical bone thickness needing to exceed trabecular thickness, the cortical border being continuous, or requiring a set amount of “background” space in the scan. Our method also provides the option to adjust the algorithm parameters or to manually adjust marrow space input to give flexibility in how the cortical - trabecular boundary in specimens with high porosity is designated. Such bespoke adjustments also allow our method to work on challenging morphologies such as the growing mouse epiphysis, where sinus volume is large relative to total volume. The flexibility of our manual pre-processing steps enables the method to work on a range of morphologies and accommodates adjustments for excluding or including features such as mineralized ligaments, and not-enclosed trabecular surfaces or presence of cortical primary osteons at the early infilling stages where most methods struggle to pinpoint these as cortex. We also reveal that measurement of bone parameters including trabecular BV/TV and anisotropy, is mostly similar between sinus-cleaned (more detailed) and non-sinus cleaned segmentation. We hope that users can replicate the methods we have specified herein in other, specifically open access software.

## 6. Access to Algorithm and CT Data

The Avizo recipe and instructions for implementation are available on our Github repository: https://github.com/evaherbst/Trabecular_Segmentation_Avizo. If you use this recipe, please cite this paper in any resulting publications. Scan data is available on our Figshare database: https://figshare.com/projects/Trabecular_and_Cortical_Bone_Segmentation_Method/99434. If you use the scans, please cite the Figshare datasets.

## 7. Acknowledgements and Funding

We thank the OA Tech+ Network (grant EP/N027264/1) the Anatomical Society (grant SSD 011018SEAL – v1-011217) for funding this project. We thank Alessandro Borghi and Great Ormond Street Hospital Charity Clinical Research Starter Grant (grant 17DD46) for the human cranial bone scan and Chaozong Liu and the European Commission via H2020-MSCA-RISE programme (BAMOS, grant 734156) and Rosetrees Trust (grant A1184) for the human femur scan. We also thank Kamel Madi for sharing his expertise in Avizo. We thank the Avizo software team, especially Julien Roussel and Alejandra Sanchez-Erostegui, for providing us home office licenses during the COVID pandemic.

## 8. Author Contributions

Eva C. Herbst designed the study, wrote the manuscript, and made the figures. Eva C. Herbst, Alessandro A. Felder and Lucinda A. E. Evans segmented the scans. Alessandro A. Felder consulted on the method development. Lucinda A. E. Evans and Andrew A. Pitsillides contributed to study design. Sara Ajami proided scans for the human femoral head and parietal bone. Behzad Javaheri provided scans for the STR/Ort and CBA mice. Eva C. Herbst, Alessandro A. Felder, Lucinda A. E. Evans, Sara Ajami, Behzad Javaheri, and Andrew A. Pitsillides all edited and revised the manuscript and approve submission of this manuscript.

## 9. Supplementary Information

Video of 3D rendered trabecular segmentation and slice data of 20 week CBA mouse epiphysis: https://figshare.com/articles/media/Video_demonstrating_trabecular_segmentation_of_a_mouse_epiphysis/14137964

